# Repurposing melanoma chemotherapy to activate inflammasomes in treatment of BRAF/MAPK inhibitor resistant melanoma

**DOI:** 10.1101/2021.08.23.457344

**Authors:** Farzana Ahmed, Hsin-Yi Tseng, Antonio Ahn, Dilini Gunatilake, Sara Alavi, Michael Eccles, Helen Rizos, Stuart Gallagher, Jessamy Tiffen, Peter Hersey, Abdullah Al Emran

**Affiliations:** Melanoma Immunology and Oncology Group, The Centenary Institute, University of Sydney, Camperdown, NSW, Australia; Melanoma Institute Australia, Crows Nest, Sydney, NSW, Australia; Central Clinical School, The University of Sydney, Camperdown, NSW, Australia; Peter MacCallum Cancer Centre, Melbourne, Victoria, Australia; Sir Peter MacCallum Department of Oncology, University of Melbourne, Parkville, Victoria, Australia; Department of Pathology, Dunedin School of Medicine, University of Otago, 270 Great King Street, Dunedin 9054, New Zealand; Maurice Wilkins Centre for Molecular Biodiscovery, Level 2, 3A Symonds Street, Auckland, New Zealand; Department of Biomedical Sciences, Faculty of Medicine, Health and Human Sciences, Macquarie University; Cutaneous Biology Research Center, Department of Dermatology, Massachusetts General Hospital, Harvard Medical School, Charlestown, Massachusetts

**Author notes:** To whom correspondence should be addressed: Peter Hersey. Joint first author (Farzana Ahmed, Hsin-Yi Tseng). Joint senior author (Peter Hersey, Abdullah Al Emran).

**Keywords:** Melanoma, resistance, inflammasome, pyroptosis, chemotherapy

## Abstract

The development of resistance to treatments of melanoma is commonly associated with upregulation of the MAPK pathway and development of an undifferentiated state. Prior studies have suggested that melanoma with these resistance characteristics may be susceptible to innate death mechanisms such as pyroptosis triggered by activation of inflammasomes. In the present studies we have taken cell lines from patients before and after development of resistance to BRAF V600 inhibitors and exposed the resistant melanoma to temozolomide (a commonly used chemotherapy) with and without chloroquine to inhibit autophagy. It was found that melanoma with an inflammatory undifferentiated state appeared susceptible to this combination when tested *in vitro* and *in vivo* against xenografts in NSG mice. Translation of the latter results into patients would promise durable responses in patients treated by the combination. The inflammasome and death mechanism involved appeared to vary between melanoma and involved either AIM2, NLRP3 or NLRC4 inflammasomes and gasdermin D or E. These preliminary studies have raised questions as to the selectivity for different inflammasomes in different melanoma and their selective targeting by chemotherapy. They also question whether the inflammatory state of melanoma may be used as biomarkers to select patients for inflammasome targeted therapy.

## Introduction

Selective inhibitors of *BRAFV600* mutations in the MAPK signal pathway and immune checkpoint blockade (ICB) ushered in a new era in melanoma treatment (Luke et al., 2017). Despite these advances, not all patients respond to such treatments and even in responding patients, therapy resistance occurs in the majority. Multiple causes of resistance to immunotherapy have been defined (Ribas and Wolchok, 2018, Sharma et al., 2017, Utzschneider et al., 2016) including defects in antigen presentation (Sade-Feldman et al., 2017) and low T cell infiltration into tumors (Li et al., 2019) whereas in targeted therapies reactivation of the MAPK pathway (Long et al., 2014, Song et al., 2017) is the commonest cause of treatment failure (Kozar et al., 2019). Adaptive resistance associated with dedifferentiation of melanoma has been described against both forms of treatment (Bai et al., 2019, Shaffer et al., 2017).

Treatments offered to patients failing ICB treatments include retreatment with anti-PD1 plus or minus anti-CTLA4 with variable success (Betof Warner et al., 2020, Olson et al., 2021). Very quickly however patients can exhaust standard treatment options leaving open the question whether chemotherapy that was commonly used prior to targeted therapies and ICB may have a role in such patients. This was examined in studies on 60 patients who had failed anti-PD1. Thirty-three patients treated with a combination of chemotherapy and ICB had a median overall survival (OS) of 3.5 years whereas patients receiving ICB alone (9) or chemotherapy alone (18) had median OS of 1.8 years. Overall response rates where 59% vs 15% in the 2 groups (Vera Aguilera et al., 2020). These studies plus anecdotal experience of responses to chemotherapy after treatment with ICB (Swami et al., 2019) suggest that chemotherapy deserves further investigation in patients who have exhausted standard treatment options.

Renewed interest in chemotherapy has also been stimulated by discovery of innate death mechanisms that can be activated by chemotherapy (Emran et al., 2020). These include ferroptosis that results from iron dependent lipoxygenase enzyme peroxidation of polyunsaturated fatty acids in cell membranes, necroptosis that is induced by death receptor ligands in cells with low or absent caspase-8 levels and pyroptosis that results from activation of inflammasomes and release of gasdermins (GSDMs). The latter cause cell death by creating pores in cell membranes resulting in loss of cell contents and release of cytokines such as IL-1β and IL-18 (Broz et al., 2020). Of particular interest were studies showing that some dedifferentiated resistant melanoma were sensitive to innate death mechanisms such as ferroptosis (Tsoi et al., 2018). GSDMs have attracted much interest as previous studies have shown that chemotherapy can activate GSDMs (Wang et al., 2017). GSDME in particular has been activated by chemotherapy agents such as Cisplatin (Zhang et al., 2019) and Doxorubicin (Yu et al., 2019). Our previous analysis of data from the cancer genome atlas (TCGA) showed that high levels of inflammasomes were associated with improved survival of patients with melanoma (Emran et al., 2020).

In the present study, most attention has been given to temozolomide (TMZ) as it is the most widely used chemotherapy against melanoma. It is a non-classic alkylating agent that is converted in the liver to its active agent-methyl-triazeno imidazole carboxamide (MTIC) (Mhaidat et al., 2007)(Becker et al., 2000). Many other drugs have been tried against melanoma but the combination of Carboplatin and paclitaxel has been the main alternative to TMZ (Flaherty et al., 2013, Hauschild et al., 2009). To maximize the effects of chemotherapy, we have included chloroquine (CQ) as an inhibitor of autophagy. The latter is a critical regulator of inflammasome activation by removal of endogenous signals that would otherwise activate inflammasomes. CQ is a known inhibitor of autophagy and several derivatives are currently in clinical trials with chemotherapy (Amaravadi et al., 2019, Rebecca et al., 2019). In the following studies we have focused particularly on whether the death mechanisms activated by the chemotherapy/CQ combination were active in BRAF/MEK resistant melanoma lines, particularly as the MEK pathway is known to regulate autophagy (Wang et al., 2009).

## Results

### *BRAFV600E* mutated melanoma lines that are resistant to dabrafenib can be killed by Temozolomide in the presence of Chloroquine

We studied the ability of dabrafenib to induce lytic cell death on matched melanoma cell lines established from patients prior to BRAF-inhibitor(i) treatment (pre) or following progression on BRAFi treatment (post). Melanoma lines established pretreatment showed variable sensitivity to treatment with dabrafenib but all the post-treatment lines were resistant to dabrafenib in LDH release assays (Figure 1a).

**Figure 1:**
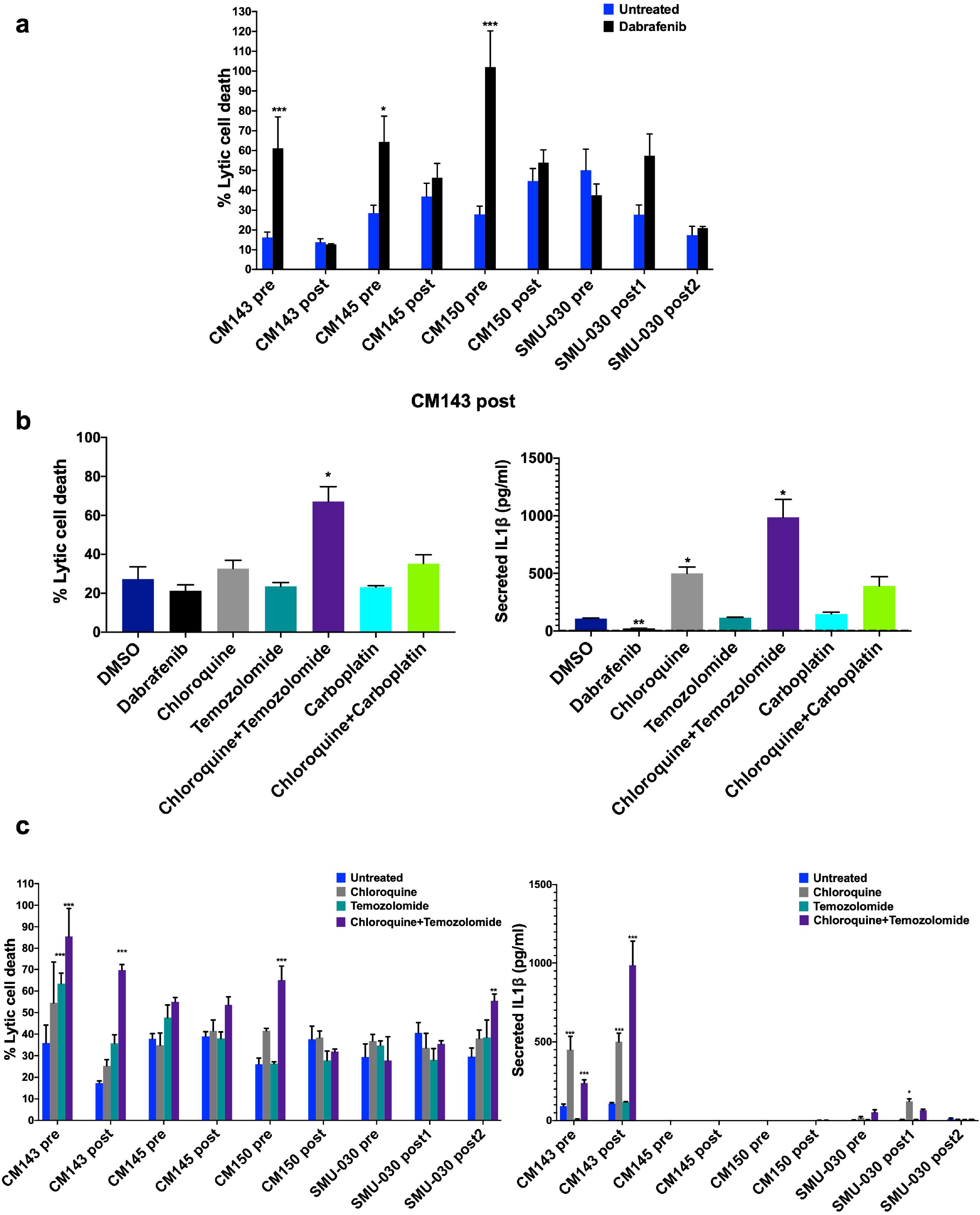
TMZ and CQ combination induced lytic cell death in the selected match-paired patient lines. **(a)** Lytic cell death was determined by the lactate dehydrogenase (LDH) release assay in the indicated patient match-paired cell lines. Cells were treated with either DMSO or dabrafenib (100nM) for 72 hours followed by LDH assay for lytic cell death (n=3). Unpaired t-test was performed to assess the significance where \**P*<0.05, \*\*\**P*<0.001. **(b)** Lytic cell death and IL1β secretion measured LDH and Cytometric Bead Array (CBA) in CM143 post cells with the indicated drugs. Cells were treated with DMSO, dabrafenib (100nM), pretreatment with chloroquine (20μM) for 1h followed by temozolomide (100μM), carboplatin (1.34μM) and pretreatment with chloroquine (20μM) for 1h followed by carboplatin (1.34μM) for 72 hours (n=3). Unpaired t-test refers to significance where \**P*<0.05, \*\**P*<0.01. **(c)** LDH assay (left panel) to assess the lytic cell death in the indicated cell lines. Cells were DMSO, pretreatment with CQ (20μM) for 1h followed by TMZ (100μM) for 72 hours. Same samples were assessed for IL-1β by CBA assay (right panel) (n=2). Multiple t-test was performed to assess the significance where \**P*<0.05, \*\**P*<0.01, \*\*\**P*<0.001.

The results of lytic cell death and secreted IL-1β in the CM143 post cell line before and after treatment with temozolomide (TMZ) or carboplatin and with or without chloroquine (CQ) is illustrated in figure 1b. The combination of agents but not the single agents induced strong lytic cell death.

When these studies were carried out in a larger panel of paired cell lines the TMZ-CQ combination increased LDH release in CM145 and SMU-030 post2 lines but not CM150 post lines (Figure 1c, left panel). There was a marked increase in IL-1β in CM143 post line but only very low release from the SMU-030 lines (Figure 1c, right panel). Stimulation with IL-1α and LPS also showed similar results in CM143 post line (Supplementary Figure S1a,b). To assess whether IL-1β could be a potential biomarker of inflammatory cell death we investigated the RNA-seq data of TCGA melanoma cohort which showed that a subset of melanoma patients with high level of IL-1β had a significant better survival compared to low patient group (Supplementary Figure S1c). Additionally, paired pre-treatment and progression tumor transcriptome data (GSE50509)(Rizos et al., 2014) from patients who progressed on BRAF inhibitor treatment had low levels of IL-1β although not significant. This suggests that level of IL-1β could be a potential biomarker of selecting patients who are likely to respond to TMZ-CQ combination (Supplementary Figure S1d).

### Upregulation of inflammatory and immune regulated pathways and downregulation of differentiation related pathways were prominent in the sensitive CM143 lines

To assess the pathways distinct in TMZ-CQ sensitive CM143 lines compared to resistant CM145 and CM150 we performed a GSEA analysis based on the RNA-seq data of the match-paired lines. This revealed upregulation of several immune associated pathways. (Figure 2a). Hallmark TNFα mediated NF-KB signaling, IFNα response and Inflammatory response were the top three upregulated pathways in the sensitive CM143 lines compared to resistant CM145 and CM150 lines (Normalized enrichment score >1.5 and p<0.001, Figure 2b). Heatmaps of the individual genes in Inflammatory response, TNFα signaling and IFNα response suggest that they were consistently upregulated in the CM143 lines compared to the CM145 and CM150 lines (Figure 2c, Supplementary Figure S2a). Additionally, several differentiation related pathways were downregulated in the sensitive CM143 lines (Supplementary Figures S2b). As shown in supplementary figure S3 there was a range of mRNA for inflammasome sensors and downstream components in the match-paired lines.

**Figure 2:**
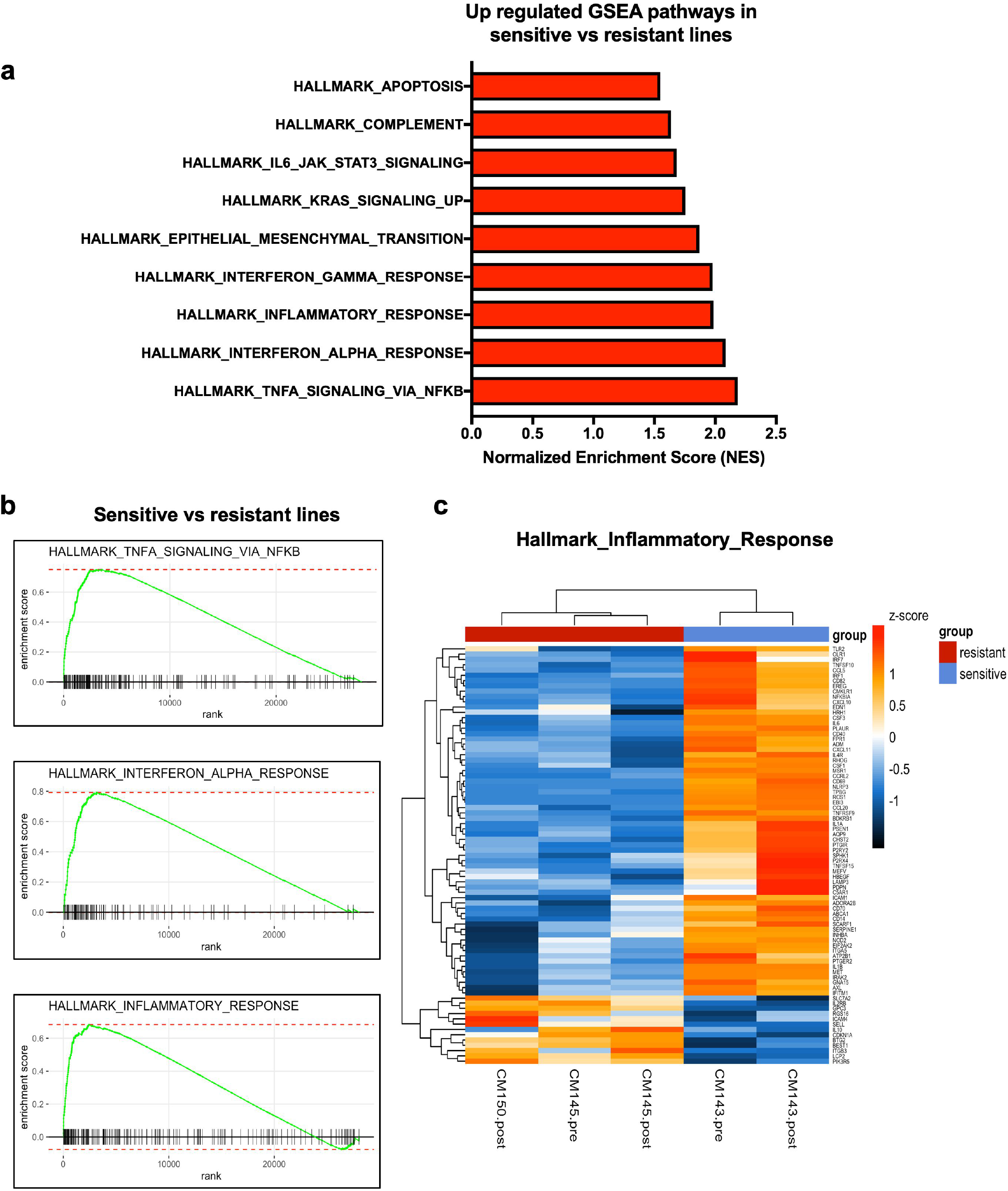
GSEA of differentially expressed genes in patient match-paired cell lines. **(a)** A forest plot showing all the upregulated pathways by GSEA analysis in the sensitive CM143 lines compared to resistant CM145 and CM150 lines. **(b)** GSEA graphs showing the top three up regulated pathways. **(c)** A heatmap showing differentially expressed genes of the individual cell lines. Z score represent the differentially expressed genes in the hallmark of inflammatory response.

### BRAFi resistant melanoma lines can be killed by activators of AIM2 and NLRP3 inflammasomes but not by activation of NLRP1

To determine which activators of inflammasomes may induce cell death in melanoma we tested not only the TMZ-CQ combination but also the known activators of NLRP3 (Nigericin), AIM2 (poly(dA:dT)), NLRP1 (Val boro-Pro (VbP) - inhibitor of DPP8/9) (Gai et al., 2019). Activators of NLRP3 and AIM2 induced lytic cell death in CM143 cells (Figure 3a). Dabrafenib did not induce any lytic cell death which confirms the result of Figure 1a. VbP was also unable to kill melanoma but was able to kill OCI-AML2 cells (Data not shown). Both IL-1α or LPS showed similar results with unstimulated condition suggesting that priming is not required to induce inflammatory cell death in CM143 lines (Figure 3a).

**Figure 3:**
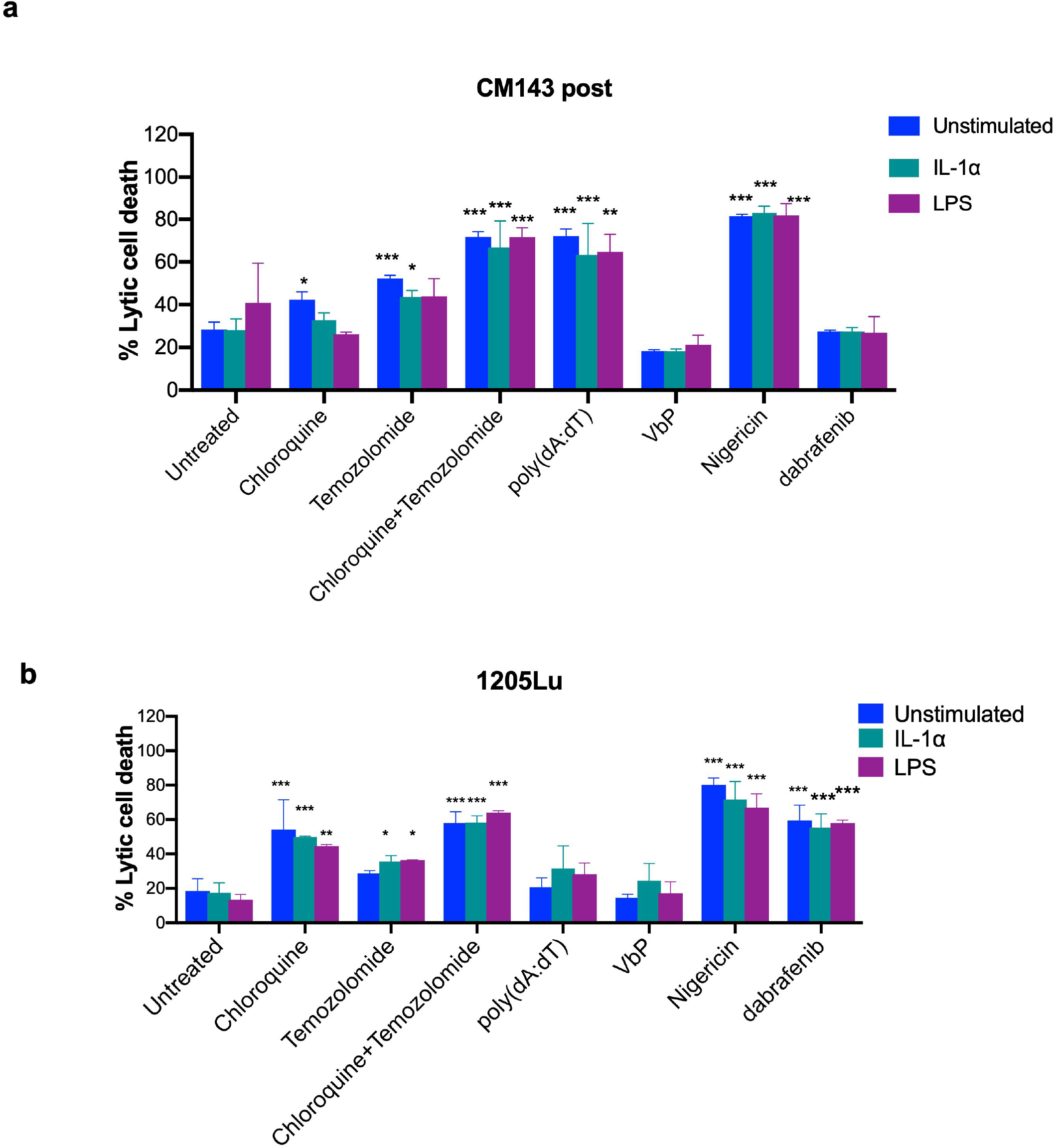
Lytic cell death with activators of inflammasomes in CM143 post and 1205Lu. CM143 post **(a)** and 1205Lu **(b)** cells were primed or unprimed with IL-1α (10ng/ml) or LPS (1μg/ml) for 24 hours and washed out. All cells were then pre-treated with chloroquine (20μM) for 1h followed by the indicated treatment for 3 days (n=2). Temozolomide (100μM); poly(dA:dT) (1μg/ml) - transfected with LyoVec; VbP (20μM); Nigericin (10μM); dabrafenib (100nM).

A similar study was carried out on the 1205Lu line described elsewhere (Okamoto et al., 2010) (Figure 3b). Similar to CM143 post, this line also did not need priming by IL-1α or LPS for killing by activation of inflammasomes. The main differences between these two cell lines were that there was no killing by activation of AIM2 by poly(dA:dT) but there was killing by dabrafenib most probably due to inhibition of BRAF V600 pathway in 1205Lu. This may possibly involve indirect activation of NLRC4 as reported by others (Hajek, Krebs et al. 2018) but has not been further investigated in these studies.

### Cell death mechanisms activated by TMZ-CQ differs between cell lines

To determine the cell death mechanism that was involved in the TMZ-CQ combination effects we used inhibitors of caspases, such as caspase-1 inhibitor (VX-765), caspase-3 inhibitor (z-DEVD-FMK) and the pan-caspase inhibitor (Q-VD-OPH). MCC950 was used to inhibit NLRP3 and disulfiram to inhibit GSDMD (Hu et al., 2020). LDH release assay suggest that TMZ-CQ mediated killing of CM143 post can be inhibited by the pan-caspase and caspase-3 inhibitor but not by the caspase-1 inhibitor (Figure 4a, left panel). There was also partial inhibition by disulfiram against GSDMD. A similar pattern was seen in assays of IL-1β release except that disulfiram did not inhibit IL-1β release (Figure 4a, right panel). Necrosulfonamide was reported to be an inhibitor of GSDMD (Rathkey et al., 2018) but a repeat of the study with necrosulfonamide did not inhibit lytic death suggesting that GSDMD may not mediate cell death induced by TMZ-CQ (Data not shown).

**Figure 4:**
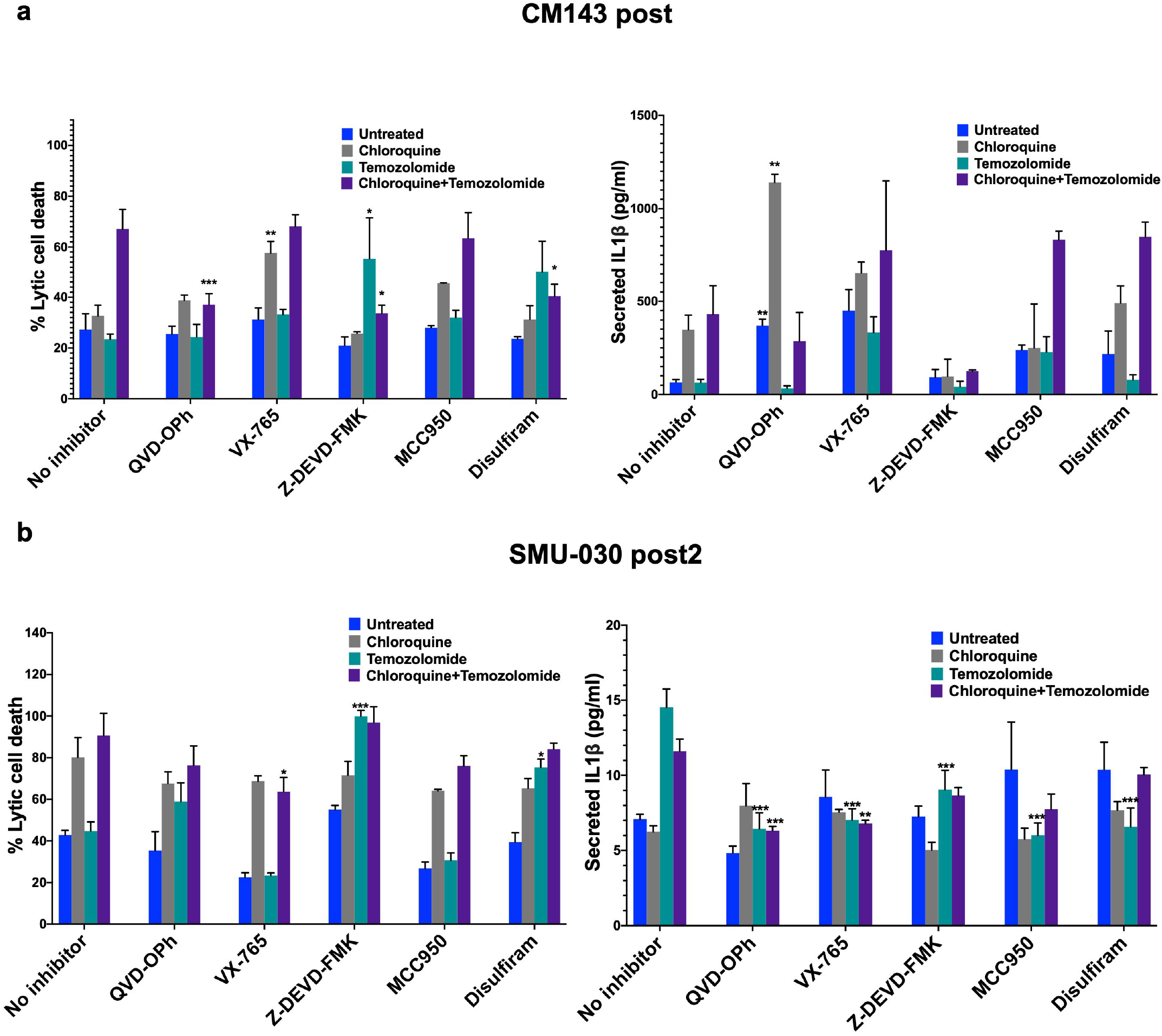
TMZ and CQ combination activated different mode of cell deaths. **(a)** CM143 post cells were pre-treated with QVD-OPH (10μM), Vx-765(10μM), Z-DEVD-FMK (50μM), MCC950(1μM) and Disulfirum (50ng/ml) for 1 hour followed by treatment with DMSO, CQ (20μM), TMZ (100μM) and combination for 72 hours. Treated cells were subjected for LDH assay for lytic cell death analysis (left panel). Supernatant from the sample samples were harvested for IL-1β by CBA (right panel) (n=2). Similar experiments were performed on SMU-030 post2 cells and analyzed by (b) LDH (left panel) and CBA assay (right panel) for lytic cell death and IL-1β respectively (n=2). Multiple t-test was performed to assess the significance where \**P*<0.05, \*\**P*<0.01, \*\*\*\**P*<0.0001. All treatments were compared to the untreated group for statistical analysis.

Similar studies were carried out against the SMU-030 post2 line (Figure 4b, left panel). Significant inhibition of IL-1β release was seen with the inhibitors of caspase 1 and pan-caspases (Figure 4b, right panel). There was also partial inhibition by MCC950 suggesting involvement of NLRP3. Similar results were seen in assays of LDH release.

### Cleavage of Gasdermins are involved in killing of BRAFi resistant melanoma by TMZ plus CQ

To further investigate the killing mechanisms, we examined the expression of sensor proteins in the CM143 cells at 24 h intervals. The sensor proteins NLRP1, NLRP3, NLRC4 and AIM2 did not undergo significant changes but cleaved caspase 3 was evident from 24 h onwards (Figure 5a). Caspase-1 cleavage was not evident which was consistent with the inability of caspase 1 inhibitors to block TMZ-CQ killing. GSDMD cleavage was evident from 24 h onwards. GSDME cleavage was also detected by 48 h (Figure 5a). These changes in caspase-3 GSDMD and GSDME were not detected in the CM142 line which did not respond to the TMZ-CQ combination (Supplementary Figure S4a, b).

**Figure 5:**
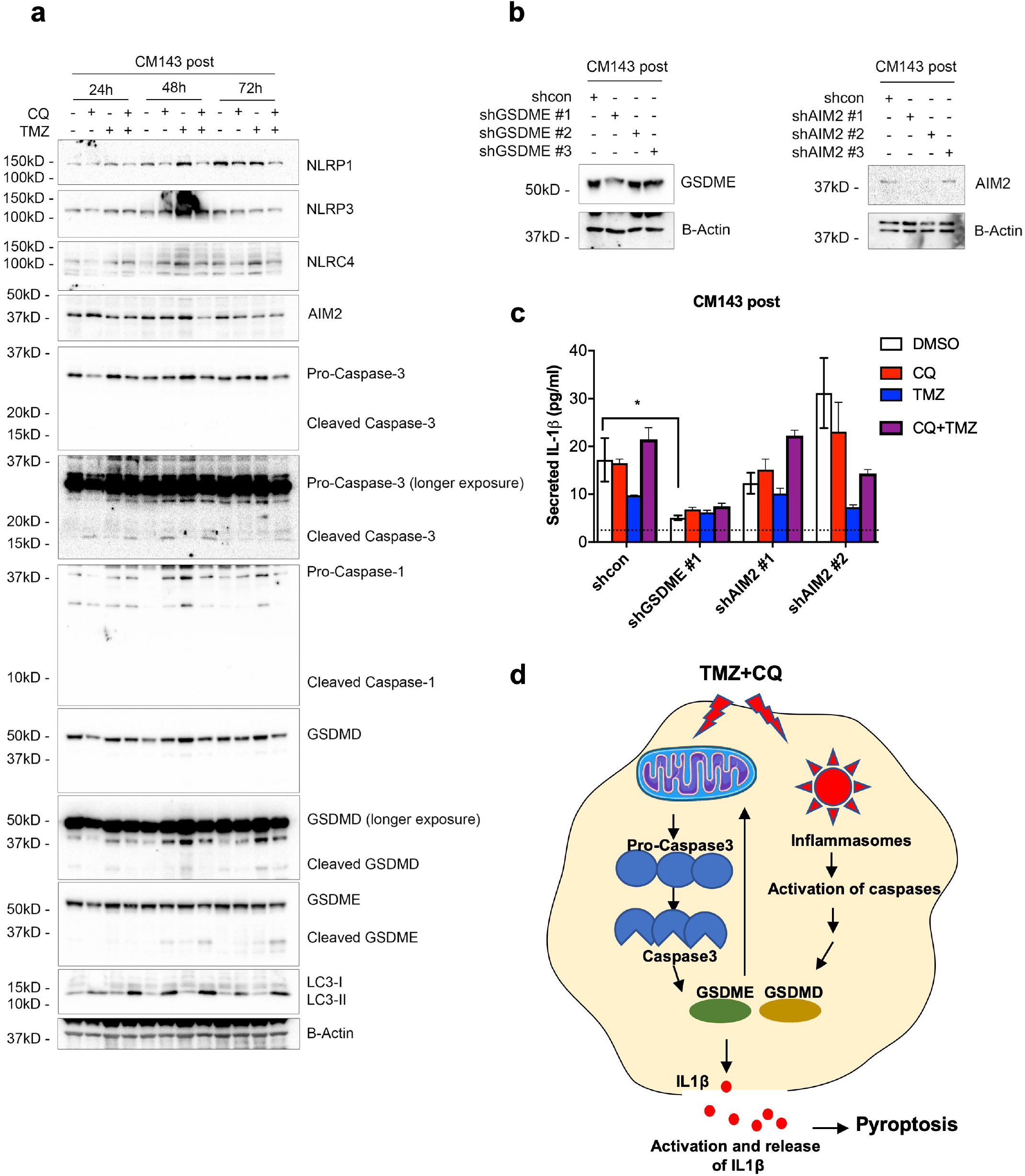
TMZ and CQ induced cleavage of Caspase-3 and Gasdermin E in CM143 post cells. **(a)** Representative western blot images of inflammasome-related proteins after the indicated treatment in CM143 post. CM143 post were pre-treated with or without CQ (20μM) for 1h followed by TMZ (100μM) for the indicated period. CQ, chloroquine; TMZ, temozolomide (n=2). **(b)** Representative western blot images of GSDME (left panel) and AIM2 (right panel) in CM143 post knockdown cells. CM143 post were transduced with the indicated constructs, expanded and sorted for positive green fluorescent cells. Lysates from the sorted cells were subjected to western blotting with GSDME or AIM2 (n=2). (**c)** CBA of secreted IL-1β in CM143 post knockdown cells. CM143 post were transduced with the indicated constructs, expanded and sorted for positive green fluorescent cells. Sorted cells were expanded and pre-treated with or without CQ (20μM) for 1h followed by TMZ (100μM) for 3 days. Supernatants collected were subjected to cytometric bead array with IL-1β beads (n=2). **(d)** A schematic illustration of proposed cell death mechanism.

Knock down of GSDME markedly inhibited IL-1β release from the CM143 cells (Figure 5b-c). Knockdown of AIM2 also reduced IL-1β but was not as marked as GSDME (Figure 5b-c). These results would be consistent with AIM2 being upstream of enzymes involved in GSDME and GSDMD cleavage whereas the cleavage of the gasdermins may also involve non AIM2 initiated events such as intrinsic apoptosis generated caspase-3 involved in cleavage of the GSDMs (Rogers et al., 2019).

We proposed a schematic representation of 2 plausible mechanisms involved in their cleavage (Figure 5d). One being inflammasome activation and cleavage by caspases like caspase-1 and -4. The other pathway being events downstream of mitochondrial permeabilization by intrinsic pathways which may include GSDME as reviewed elsewhere (Rogers et al., 2019). In this view GSDME is seen not only as an amplifier of pyroptosis but also as an initiator of mitochondrial changes similar to BH3 proapoptotic proteins.

### TMZ alone and in combination with CQ inhibits the growth of melanoma CM143 xenografts

The *in vivo* study was preceded by a drug tolerability study. Single inhibitors or the combination of 25mg/kg TMZ and CQ did not adversely affect the body weight and well tolerated in the NSG mice after 14 days of treatment (Supplementary Figure S5a-c).

CQ had minimal effects on tumor growth but TMZ alone and TMZ plus CQ markedly suppressed growth of the tumors (Figure 6a-c, Supplementary Figure S6). After two weeks of treatment mice were observed without any treatment in the TMZ and combination group. Regrowth of the tumors was evident by day 25 in the TMZ alone mice but not in the mice receiving the combination of drugs. Combination treatment reduced the tumor growth by 81% at day 32 compared to TMZ group (Figure 6d). All the treatments were tolerated in mice. The drugs either singly or in combination had no significant effects on body weight of the mice (Figure 6e).

**Figure 6:**
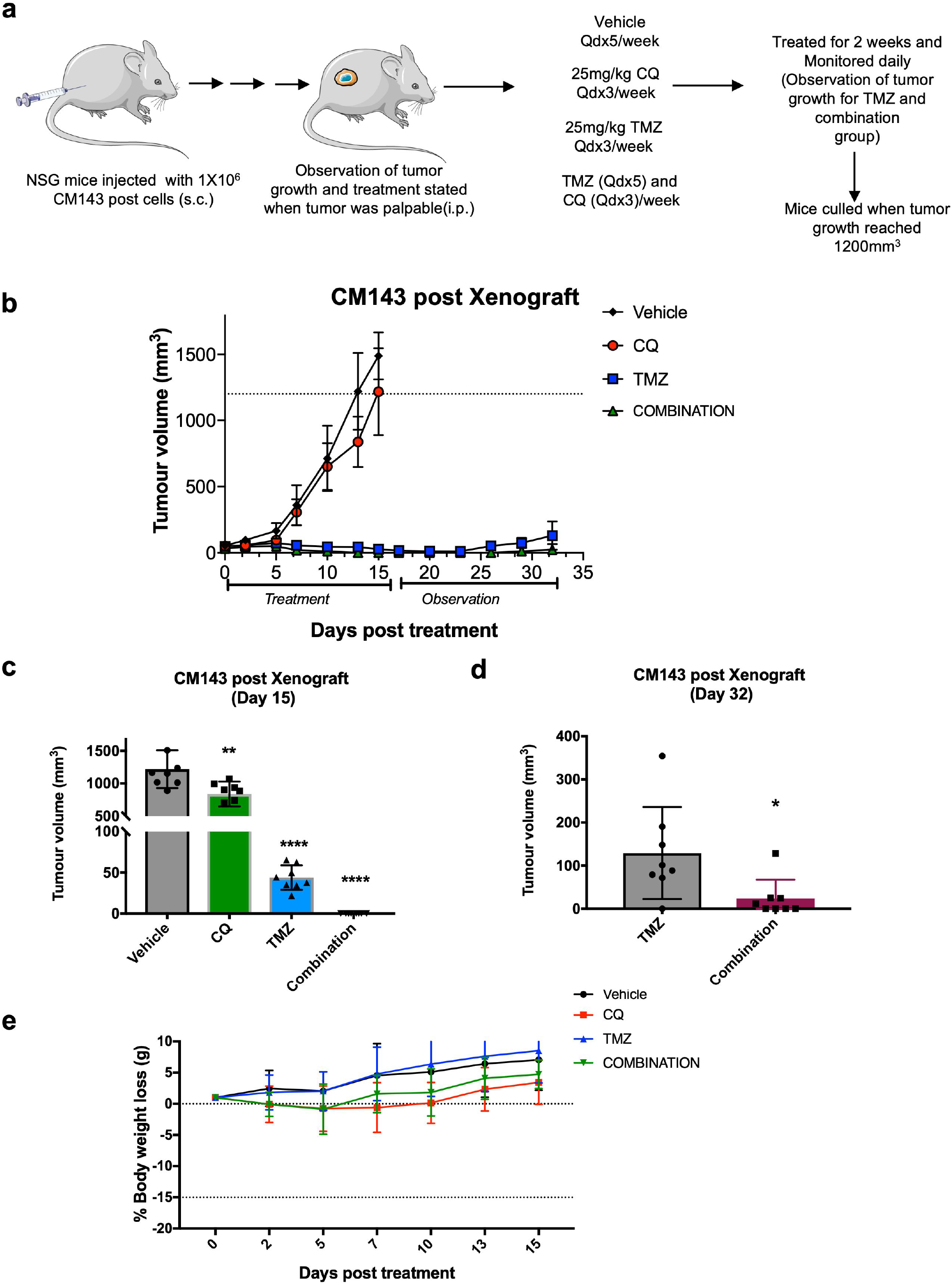
TMZ alone or combination with CQ markedly reduce tumor growth *in vivo*. **(a)** Dosing schedule of the indicated drugs. **(b)** Tumor volume of CM143 post xenograft. NSG mice were injected with CM143 cells and when tumor was palpable mice were randomized and treated with sterile PBS (vehicle), 25mg/kg TMZ, CQ and combination for two weeks (n=8). Treatments were withdrawn on day 14 and tumor growth was observed for another 18 days in the TMZ and combination group. (**c-d)** Bar plots showing the tumor volume on day 15 and 32. Unpaired t-test refers to significance where \**P*<0.05, \*\**P*<0.01, \*\*\*\**P*<0.0001. (**e)** Percentage of body weight loss in the tumor bearing treatment group. All the weights were normalised to the starting point of each treatment group.

## Discussion

The purpose of this study was to examine whether standard chemotherapeutic agents used against melanoma could kill melanoma by recruitment of innate death mechanisms and whether this could be targeted to melanoma that were resistant to targeted treatments. In the 4 pairs of cell lines studied all 4 of the resistant pair lines were resistant to treatment with dabrafenib which is a commonly used inhibitor of BRAFV600 signaling. TMZ or carboplatin had very little direct toxic effects on the cell lines *in vitro* but when tested in combination with CQ there was a significant increase of cell death in 2 of the BRAFi resistant lines measured in LDH lytic assays and or release of IL-1β. Of note, cell line CM143 was reported previously as having constitutive activation of NF-kB and high constitutive production of a range of cytokines such as IL-1β, IL-6 and IL-8 (Gallagher et al., 2014). In this respect it is similar to the late stage 1205Lu line studied by Okamoto et al which had constitutive secretion of IL-1β as well as IL-6 and IL-8 (Okamoto et al., 2010). We found that the latter line was also sensitive to the TMZ-CQ combination so that further studies are needed to determine whether an undifferentiated state and constitutive release of cytokines may be a common factor underlying the cell death response to TMZ-CQ. The MAPK pathway upregulates autophagy (Liu, Lai et al. 2014) so it is also possible that this could be part of the sensitivity to TMZ-CQ in the resistant lines with reactivated MAPK pathways (Liu et al., 2014).

The anticancer effects of TMZ is believed to be primarily methylation of guanine which results in DNA strand breaks that can induce apoptosis via p53 activation or other pathways (Kaina, 2019, Mhaidat et al., 2007, Zhang et al., 2012). Its primary action can be limited by repair enzymes such as methylguanine methyl transferase (MGMT) or base excision repair enzymes. These repair enzymes contribute to its relative lack of toxicity against normal tissues (Ranson et al., 2007). We speculated that a secondary effect of TMZ might be to induce release of double stranded DNA from the nucleus and thereby activate AIM2 by binding to the C terminal HIN domain of AIM2 (Wang et al., 2020). Other DNA sensors that activate interferon regulatory factors that induce production of type 1 interferons may also be activated. This speculation receives support from studies showing that single or double stranded DNA has been detected constitutively in the cytosol of human cancer cells and these can be increased by genotoxic agents (Shen et al., 2015).

Indeed, GSEA analysis of the match-paired lines showed a significant upregulation of TNF-α mediated NF-KB, interferon alpha and inflammatory pathways in the TMZ-CQ sensitive CM143 lines compared to resistant CM145 and CM150 lines. This supports the findings that constitutive expression of inflammatory signatures is crucial for TMZ-CQ sensitivity in the BRAFi resistant match-paired lines. We have published elsewhere (Ahn et al., 2021) that CM143 has a dedifferentiated state as defined by Tsoi et al with low MITF and SOX10.

To obtain evidence about the mechanism involved we first activated a range of inflammasomes to see which caused death in-vitro of CM143 resistant cells. This showed that activators of AIM2 or NLRP3 but not NLRP1 or NLRC4 could induce cell death. This contrasted with results on the 1205Lu line which was not killed by AIM2 activators but was killed by NLRP3 activators. The NLRP3 inflammasome did not appear to be activated by TMZ-CQ in CM143 melanoma in that TMZ induced cell death was not blocked by the MCC950 inhibitor of NLRP3 (Tapia-Abellán et al., 2019). To further explore the cell death mechanism in this line we used inhibitors of caspase-1, -3 and a pancaspase inhibitor expecting that the caspase-1 inhibitor would inhibit any inflammasome mediated killing. Instead, death of CM143 cells was blocked only by inhibitors of caspase-3 and the pancaspase inhibitor. Inhibitors of necroptosis (necrostatin-1) had minimal effects and disulfiram or necrosulfonamide against GSDMD (Hu et al., 2020, Rathkey et al., 2018) did not inhibit cell death. These results had similarities to prior studies showing that GSDME (DFNA5) could be activated by caspase-3 and that GSDME (Rogers et al., 2017) could also permeabilise mitochondria causing release of cytochrome c so resulting in cell death by both pyroptosis and apoptosis (Rogers, Fernandes-Alnemri et al. 2017). These results contrasted with studies on the SMU-030 line where inhibitors of both caspase-1 and pan-caspases as well as NLRP3 resulted in reduced killing *in vitro*. This was consistent with death induced by activation of NLRP3 as described elsewhere (Wang et al., 2017, Zhang et al., 2020). This interpretation was supported by the western blot studies showing cleavage of caspase-3 as an early event in the CM143 cells as well as cleavage of both GSDME and GSDMD. There was no evidence for cleavage of caspase-1. Knockdown of GSDME also inhibited TMZ-CQ induced IL-1β release. Knockdown of AIM2 gave partial inhibition of IL-1β release consistent with involvement of AIM2 inflammasomes.

The effectiveness of the TMZ-CQ combination shown *in vitro* received strong support from studies against the resistant line grown as xenografts in NSG mice in that although regrowth of melanoma was seen after 20 days in the TMZ alone treated group, no regrowth was documented up to 30 days of follow up in the TMZ-CQ treated group. If these results were translated to patients it would be equivalent to recurrence of melanoma after an initial response to TMZ alone compared to continuing remission in patients treated with TMZ plus chloroquine.

These studies leave unanswered several important questions about the role of inflammasomes in treatment of melanoma. Not all the resistant lines responded to the TMZ-CQ combination so one question is how melanoma that respond to TMZ-CQ may be identified. The CM143 line is representative of undifferentiated cells such as those undergoing adaptive epithelial mesenchyme resistance like states. Such states having low *SOX10* and *MITF* expression and increased inflammatory cytokine production as also shown in 1205Lu cells. The SMU-030 cells have BRAFi resistance due to MAPK pathway reactivation and may be representative of a second group of melanomas that are sensitive to NLRP3 activation but requiring inducers such as Toll ligands to increase their sensitivity. Further studies are needed on this. The CM143 line was associated with constitutive PD-L1 expression and further study is needed to see if this will be a useful marker of sensitivity to the TMZ-CQ combination. Most of the lines in the study except that from CM143 and CM142 had reactivation of MAPK (Gallagher et al., 2018).

A second question is whether activation of inflammasomes in the tumor is necessarily the best outcome as activation of inflammasomes may not always be beneficial. Eg NLRP1 in human melanoma cell lines 1205Lu and HS294T was found to be predominantly cytoplasmic and appeared to inhibit caspase 3/7 activity. Knockdown of the inflammasomes resulted in inhibition of cell growth and induction of apoptosis (Zhai et al., 2017). Proteomic studies have also shown that inflammatory proteins like C reactive proteins (CRP) are related to an adverse prognosis in melanoma (Karlsson et al., 2021).

Nevertheless our previous analysis of melanoma patient data in the TCGA showed that high levels of AIM2, NLRP3, NLRC4 and NLRP1 were associated with a good prognosis (Emran et al., 2020). The prognostic benefit of NLRP3 and NLRP1 was only seen in patients with high TIL levels suggesting the benefit was dependent on immune responses rather than intrinsic properties of the cells (Emran et al., 2020). Other preclinical studies have pointed to the importance of activation of inflammasomes in immune cells as mediators of responses against tumors. In the B16 model the protective effect of a vaccine was mediated by CD8 T cells and was dependent on activation of GSDME in the tumor by granzymes from the T cells that resulted in pyroptosis in the cancer cells. Granzyme A was also able to activate GSDME (Zhang et al., 2020). Further evidence for GSDME as a tumor suppressor was the high incidence of inactivating mutations in human cancers (Zhou et al., 2020).

In conclusion, melanoma that fail targeted therapy with BRAF inhibitors appear to be potentially susceptible to cell death resulting from activation of inflammasomes by exogenous agents like chemotherapy with TMZ in combination with chloroquine. The possible importance of this is shown by the induction of prolonged remission of highly malignant melanoma xenografts in NSG mice. Cell death in-vitro appeared dependent on inhibition of autophagy which is considered one of the survival pathways resulting from MAPK re-activation. Sensitivity may also be associated with undifferentiated melanoma that exhibit constitutive activation of inflammasomes and release of inflammatory cytokines and PD-L1 expression. Although human melanoma cells express a range of different inflammasomes the ability of individual inflammasomes to induce cell death appears cell line dependent and to vary between different melanoma. Further studies are needed to determine whether different chemotherapy agents can be tailored for particular inflammasomes. These results also point to the need for further studies using melanoma models in mice with intact immune systems to better understand the relative importance of direct effects on inflammasomes within tumors versus activation of the immune system.

## Materials and Methodology

Melanoma cell lines from patient 1, 3, 4 and 6 were established from patients entered into the Roche “BRIM2” phase II study of vemurafenib (Ribas et al., 2011). The SMU-030 pre and post lines were established before and during progression whilst on treatment with dabrafenib and trametinib. (Becker et al., 2014, Gowrishankar et al., 2012). All animal studies were performed according to Australian Code of Practice for the Care and Use of Animals for Scientific Purposes and with approval from the NSW Sydney Local Health District Animal Ethics Committee (2019/014B). Lytic cell death was assessed using lactate dehydrogenase (LDH) release assay according to manufacturer instructions (Thermofisher Fisher Scientific, Waltham, MA). IL-1β was detected using cytometric bead array (CBA) following manufacturer’s instructions (BD Biosciences, Franklin Lakes, NJ). Sample preparation, protein quantification and immunoblots were performed as described previously (Emran et al., 2021, Tseng et al., 2020). Functional assay using lentiviral packaging and transduction were done according to our previous protocol (Tseng et al., 2020). Detailed materials and methodology are provided in the supplementary file.

### Statistical analysis

All the statistical analyses were performed using GraphPad Prism software. Statistical significance was assessed by unpaired t-test, multiple t-test or two-way ANOVA test where \**P*<0.05, \*\**P*<0.01, \*\*\**P*<0.001 and \*\*\*\**P*<0.0001. Statistical significance of the survival was computed by Logrank P value where P<0.05 refers to significance.

## Data availability statement

RNA-seq data of the patient match-paired lines can be accessed through- https://www.ncbi.nlm.nih.gov/geo/query/acc.cgi?acc=GSE107622.

## Acknowledgments

The authors would like to thank the Sydney Cytometry Core Research Facility and Animal House Facility for their technical support.

## Author Contributions

Conceptualization: FA, HYT, AAE, PH. Investigation: FA, HYT, AAE, AS, DG. Result analysis: FA, HYT and AAE. Statistical analysis: FA, HYT and AAE. Original Draft preparation: FA, HYT, AAE and PH. Writing Review and Editing: FA, HYT, JT, SG, AAE and PH. Coordination and funding acquisition: AAE, JT and PH.

