## Supplementary material for "Repurposing melanoma chemotherapy to activate inflammasomes in treatment of BRAF/MAPK inhibitor resistant melanoma": Methods, table and supplementary figures

[†]Joint first author

[‡]Joint senior author

ORCID IDs:

#### Materials and Methods

##### Cell Culture

Melanoma cell lines from patient 1, 3, 4 and 6 were established from patients entered into the Roche “BRIM2” phase II study of vemurafenib (Ribas et al., 2011). The cell lines were established from tumor biopsies prior to treatment and during relapse from treatment with vemurafenib (labelled “pre” and “post” respectively). A summary of the cell line details is shown in Table 1. Their origin is referred to elsewhere (Lai et al., 2012). These studies were approved by the Hunter and New England Research Ethics Committee. The SMU0-30 pre and post lines were established before and during progression whilst on treatment with dabrafenib and trametinib (Becker et al., 2014, Gowrishankar et al., 2012). The 1205Lu lines have been described elsewhere and were originally isolated from melanoma at the Wistar Institute (Okamoto et al., 2010). Melanoma cell lines were cultured in Dulbecco’s modified Eagle medium (DMEM) containing 10% fetal calf serum (FCS) (AusGeneX, Brisbane, Australia). All cells were maintained at 37°C in 5% CO2. All human cell lines have been authenticated using STR profiling within the last 3 years and all experiments were performed with mycoplasma-free cells.

Table1: Details of the patient derived match-paired cell lines.

| Source patient | Cell line name | Treatment | Mutation |
|---|---|---|---|
| Patient 1 pre | CM143 pre | Vemurafenib | *BRAFV600E* |
| Patient 1 post | CM143 post | | *BRAFV600E* |
| Patient 3 pre | CM142 pre | Vemurafenib | *BRAFV600E* |
| Patient 3 post | CM142 post | | *BRAFV600E* |
| Patient 4 pre | CM150 pre | Vemurafenib | *BRAFV600E* |
| Patient 4 post | CM150 post | | *BRAFV600E* |
| Patient 6 pre | CM145 pre | Vemurafenib | *BRAFV600E* |
| Patient 6 post | CM145 post | | *BRAFV600E* |
| Patient 30 pre | SMU0-30 pre | Dabrafenib and Trametinib | *BRAFV600E* |
| Patient 30 post | SMU0-30 post 1, 2 | | *BRAFV600E* |

##### Antibodies and reagents

Antibodies used were purchased from Cell Signaling Technology (Danvers, MA) [NLRC4, ASC, NLRP3, AIM2]; Abcam (Cambridge, UK) [GSDMD, GSDME, NLRP1]; Sigma-Aldrich (St Louis, MO) [β-Actin (AC-74)]. The pan-caspase inhibitor, Q-VD-OPh, was purchased from SM Biochemicals (Anaheim, CA), the other caspase inhibitors (Z-DEVD-FMK and VX-765), Val-boro Pro (VbP), Chloroquine and Temozolomide were from MedChemExpress (Monmouth Junction, NJ). VbP was reconstituted in DMSO containing 0.1% TFA to prevent compound crystalisation. Recombinant human IL-1α was from Stemcell (Vancouver, BC) and LPS was from Sigma-Aldrich (St Louis, MO).

##### Lactate Dehydrogenase Cytotoxicity Assay

Lactate dehydrogenase (LDH) release was measured using the CyQUANT LDH Cytotoxicity Assay kit (Thermo Fisher Scientific, Waltham, MA) following the manufacturer's instructions. Briefly, cells were seeded into 96-well plates and treated as indicated with compounds. Supernatants were collected and cells were lysed with distilled water and freeze-thawed once. Cytotoxicity was determined by the formula below:

$$\% \text{ Cytotoxicity} = \frac{absorbance\ reading\ from\ supernatant}{(absorbance\ reading\ from\ supernatant\ and\ lysate\ mix)X2}\ X\ 100\%$$

##### IL-1β detection

IL-1β was measured using cytometric bead array (CBA) following manufacturer's instructions (BD Biosciences, Franklin Lakes, NJ). Briefly, cells were seeded into 96-well plates and treated as indicated with compounds. Supernatants were collected for measuring extracellular IL-1β. Samples were incubated with CBA IL-1β beads, followed by a PE-tagged secondary detection antibody. Samples were acquired by FACSCanto II (BD Biosciences, Franklin Lakes, NJ) and analysed by flowjo.

##### Western blotting

Western blot analysis was carried out as described previously (Tseng et al., 2020). Labelled bands were detected by Clarity horseradish peroxidase chemiluminescence kit (Bio-Rad, Gladesville, NSW), and images were captured with the ChemidocMP image system (Bio-Rad, Gladesville, NSW).

**Lentiviral production and transduction**

Lentiviral vectors pSIH-H1-copGFP was received as a gift from Prof Rizos (Macquarie University, North Ryde). Lentiviral vectors pSIH-H1-copGFP-shAIM2 #1, pSIH-H1-copGFP-shAIM2 #2, pSIH-H1-copGFP-shAIM2 #3, pSIH-H1-copGFP-shGSDME #1, pSIH-H1-copGFP-shGSDME #2 and pSIH-H1-copGFP-shGSDME #3 were constructed in-house (Supplementary Table S1). Vectors were transfected into HEK293T packaging cells using the calcium phosphate precipitation method as described previously (Tseng et al., 2020). Cells were transduced with virus in the presence of polybrene (8μg/ml) (Sigma-Aldrich, Castle Hill, NSW). Cells positive for green fluorescence were sorted by FACSMelody (BD Biosciences, North Ryde, Australia).

**Tumor formation and treatment *in vivo***

All animal experiments were performed in accordance with the Australian Code of Practice for the Care and Use of Animals for Scientific Purposes and with approval from the NSW Sydney Local Health District Animal Ethics Committee (2019/014b). The xenograft study was preceded by a toxicity study. Previously it has been shown that NOD SCIDγ mice can tolerate up to 25mg/kg CQ and 25mg/kg TMZ dosage as a single agent. For the drug tolerability study, 6-8 weeks old male mice were randomized into three groups (n=3). Mice were treated with carrier control; sterile PBS, CQ 25mg/kg and 25 mg/kg TMZ (qdx5 per week, i.p.) and combination with CQ 25mg/kg and 25 mg/kg TMZ (qdx5 per week, i.p.) for 14 days. Mice weight was monitored daily. $2x10^6$ CM143 post melanoma cells were subcutaneously injected into the right flank NOD SCIDγ female mice aged 6-8 weeks to form xenograft tumors. Once tumors were palpable (~$50mm^3$), mice were randomized into 4 different groups consisting of 8 mice in each group. Mice were treated with either carrier control- sterile PBS or 25mg/kg CQ (Lakhter et al., 2013) (qdx3 per week, i.p.), 25mg/kg TMZ (qdx5 per week, i.p.)(Cao et al., 2019) or in combination (qdx3 per week, i.p.) for 14 days. Tumors growth were monitored for another three weeks until the tumors volume reached maximum ethical end point of $1200mm^3$. Tumors and weights were measured thrice a week. Mice were culled if the tumor volume ≥ $1200mm^3$ or weight loss ≥ 15% or other signs of illness/distress.

**RNA-seq and GSEA analysis:**

RNA-seq data was obtained from GSE107622 (Chatterjee et al., 2018). Adaptors were trimmed using cleanadaptors (Stockwell et al., 2014). The FASTQ reads were pesduo-aligned to the hg38 reference genome using Kallisto (version 0.44.0) (Bray et al., 2016). Genomic annotations were obtained from GENECODE (Release 28, GRCh38.p12) and the gene-level count data were imported into R using Tximport (Soneson et al., 2015). Differential expression analysis was performed using DESeq2 (Love et al., 2014). Gene set enrichment analysis (GSEA) was performed using the fgsea R package (version 1.14.0). To generate the heatmaps, the raw count values were normalised using size factors generated from the DESeq2 package and subsequently z-scores were calculated. The pheatmap package (https://CRAN.R-project.org/package=pheatmap) was used to generate all the heatmaps.

**Survival analysis:**

Expression data of the skin cutaneous melanoma (SKCM) was retrieved from TCGA data. Kaplan-Meier plots were generated in Graphpad Prism 8 using overall survival and RNA-seq RSEM data SKCM cohort (Emran et al., 2021). Patients were dichotomised into groups above and below the median value of the respective gene. Statistical significance was determined by logrank p-value.

#### Supplementary S1

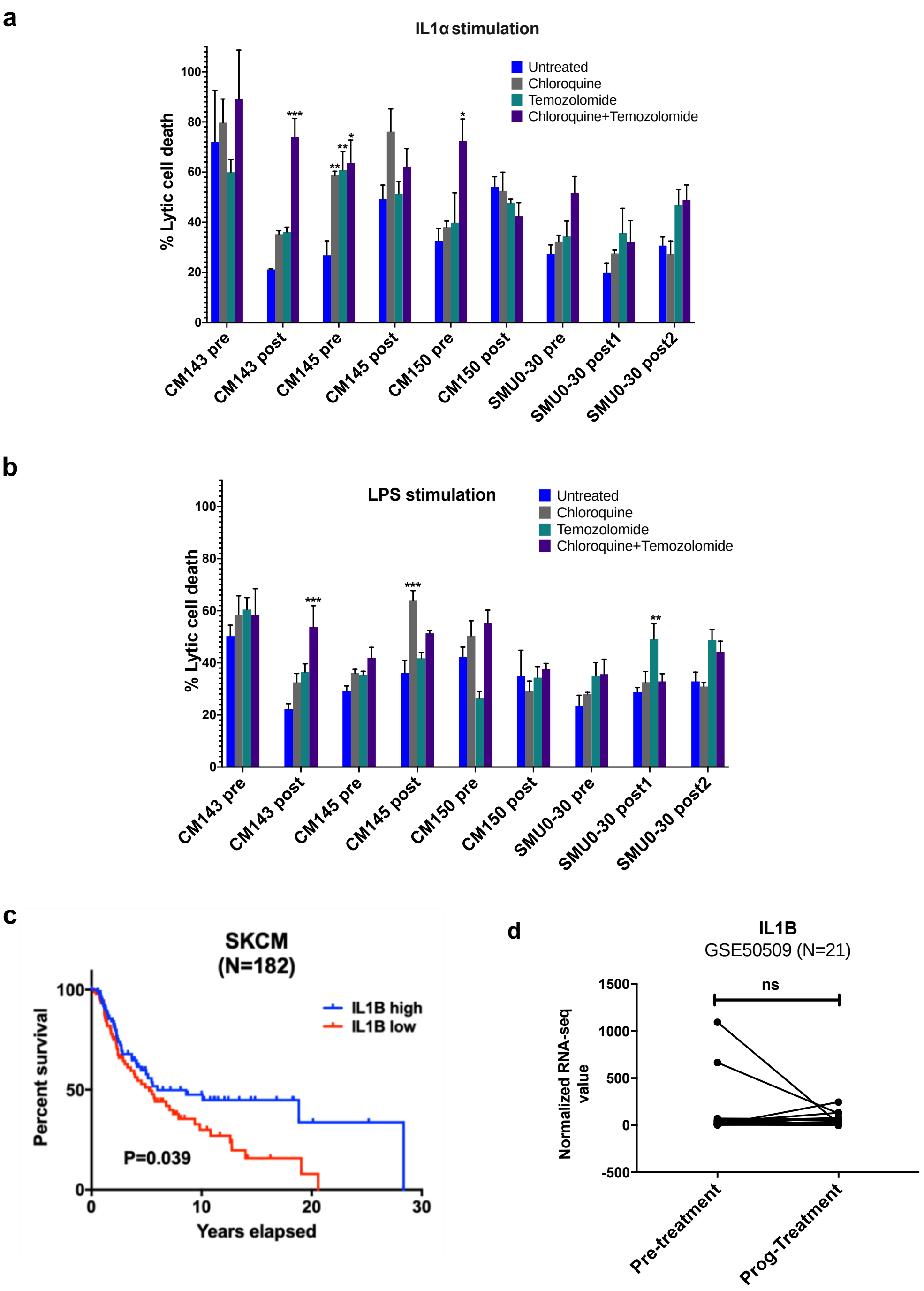


**Supplementary S1: Lytic cell death analysis with different stimulation, survival and match-paired expression analysis of IL-1β.** Bar graphs showing the lytic cell death analysis with (**a**) IL-1α and (**b**) LPS stimulation upon treated with the indicated drugs for 72 hours. Statistical analysis was performed by multiple t-test where **P*<0.05, ***P*<0.01, ****P*<0.001. **(c)** A KM plot showing the survival of subset of skin cutaneous melanoma (SKCM) patients based on median expression of IL1β. Logrank P<0.05 refers to significance. (**d**) Expression of IL1β in match paired BRAFi resistant patient derived from GSE50509 (Rizos et al., 2014).

#### Supplementary S2

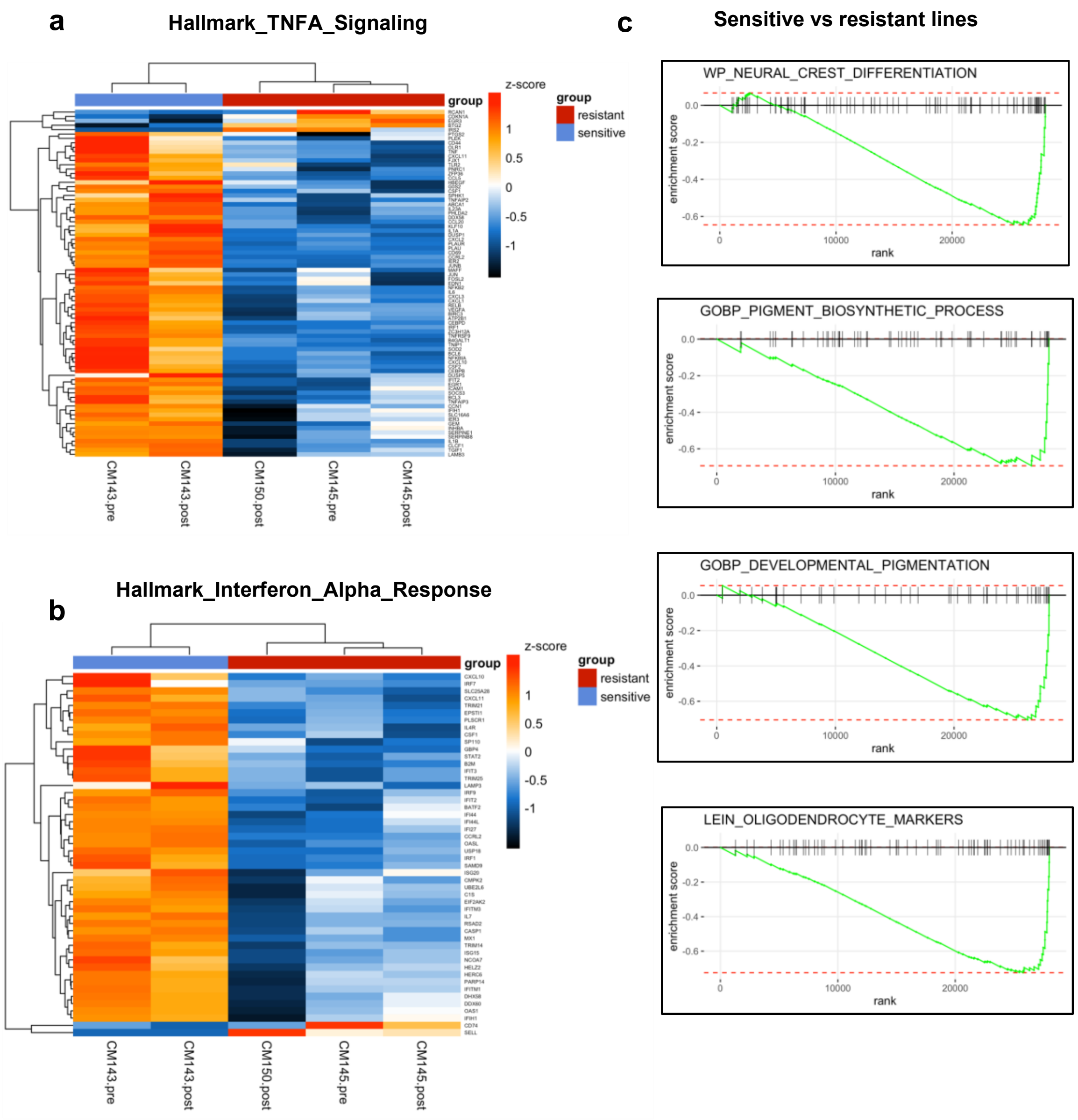


**Supplementary S2: Heatmaps and GSEA plots of differentially expressed genes in patient match-paired lines.** **(a)** Heatmaps showing differentially expressed genes of individual cell lines for hallmark TNFA signalling and interferon alpha response pathways. Z-score refers to the significant differential expressed genes. **(b)** GSEA plots for downregulated pathways in the sensitive CM143 lines compared to resistant CM145 and CM150 lines.

#### Supplementary S3

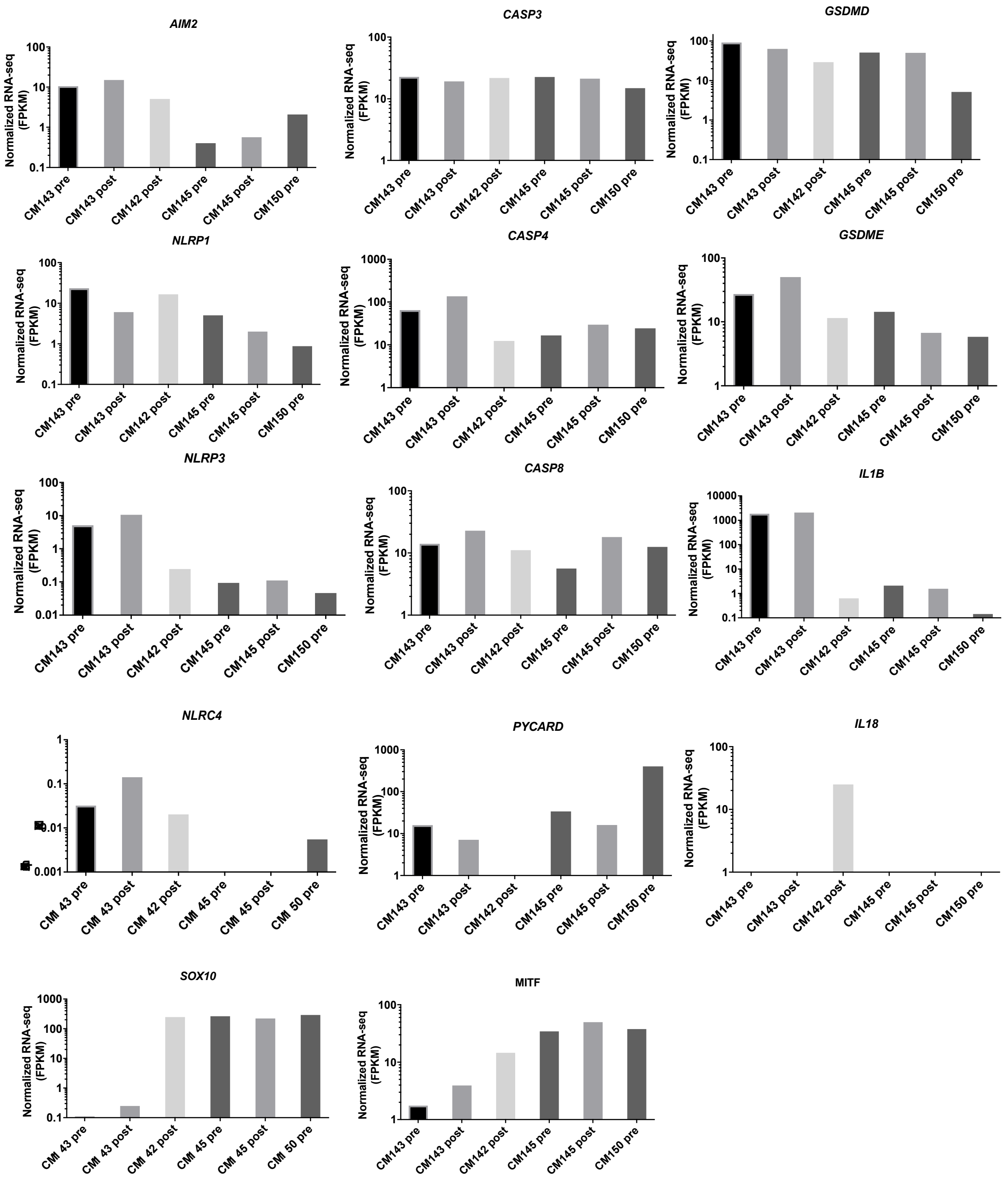


**Supplementary S3: Differential gene expression of inflammasomes and caspase family genes.** Bar graphs showing the indicated gene expression value in the CM lines.

#### Supplementary S4

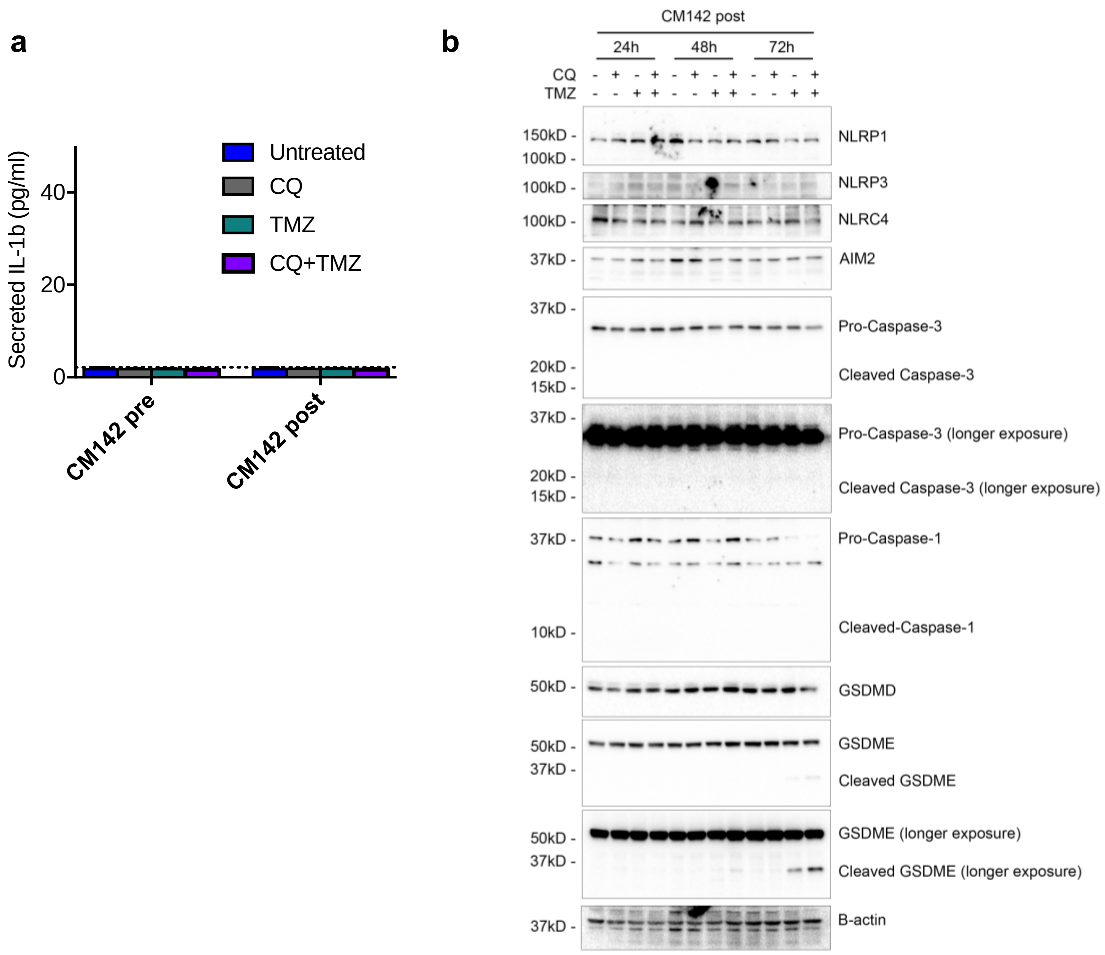


**Supplementary S4: CQ and TMZ induced effect on inflammasomes and caspase family proteins. (a)** CBA assay in CM142 pre and post cells with the indicated drugs. **(b)** Representative western blot images of inflammasome-related proteins after the indicated treatment in CM142 post. CM142 post were pre-treated with or without CQ (20µM) for 1h followed by TMZ (100µM) for the indicated period. CQ, chloroquine; TMZ, temozolomide

### Supplementary S5

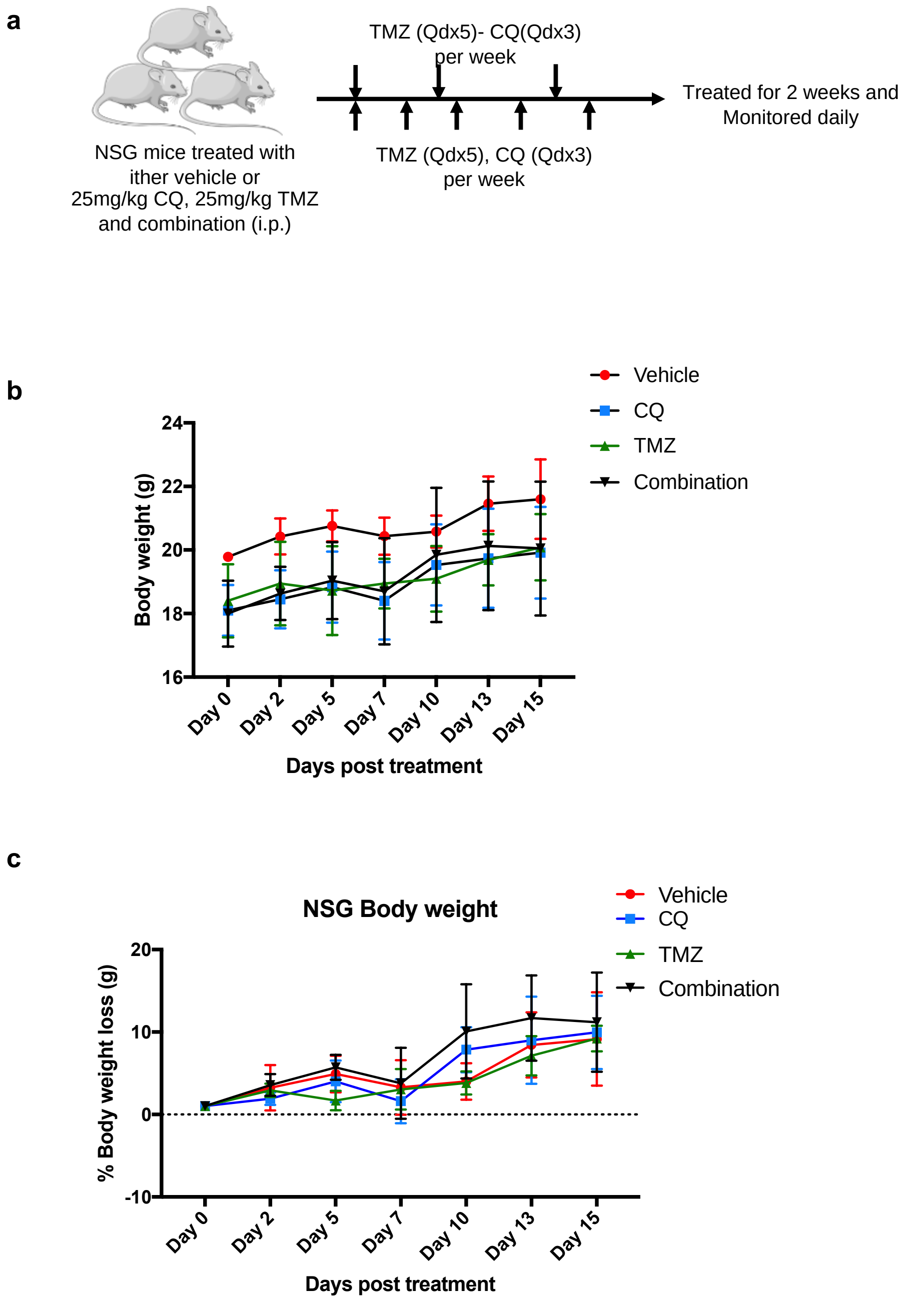


**Supplementary S5: Drug tolerability study on non-tumour bearing NSG mice(a)** Dosing schedule of the indicated drugs for tolerability study (n=3). **(b)** Mean body weight of the individual treatment group. **(c)** Percentage of body weight changes in the treatment. Weights were normalized to the starting point of each treatment group.

### Supplementary S6

**CM143 post Xenograft**

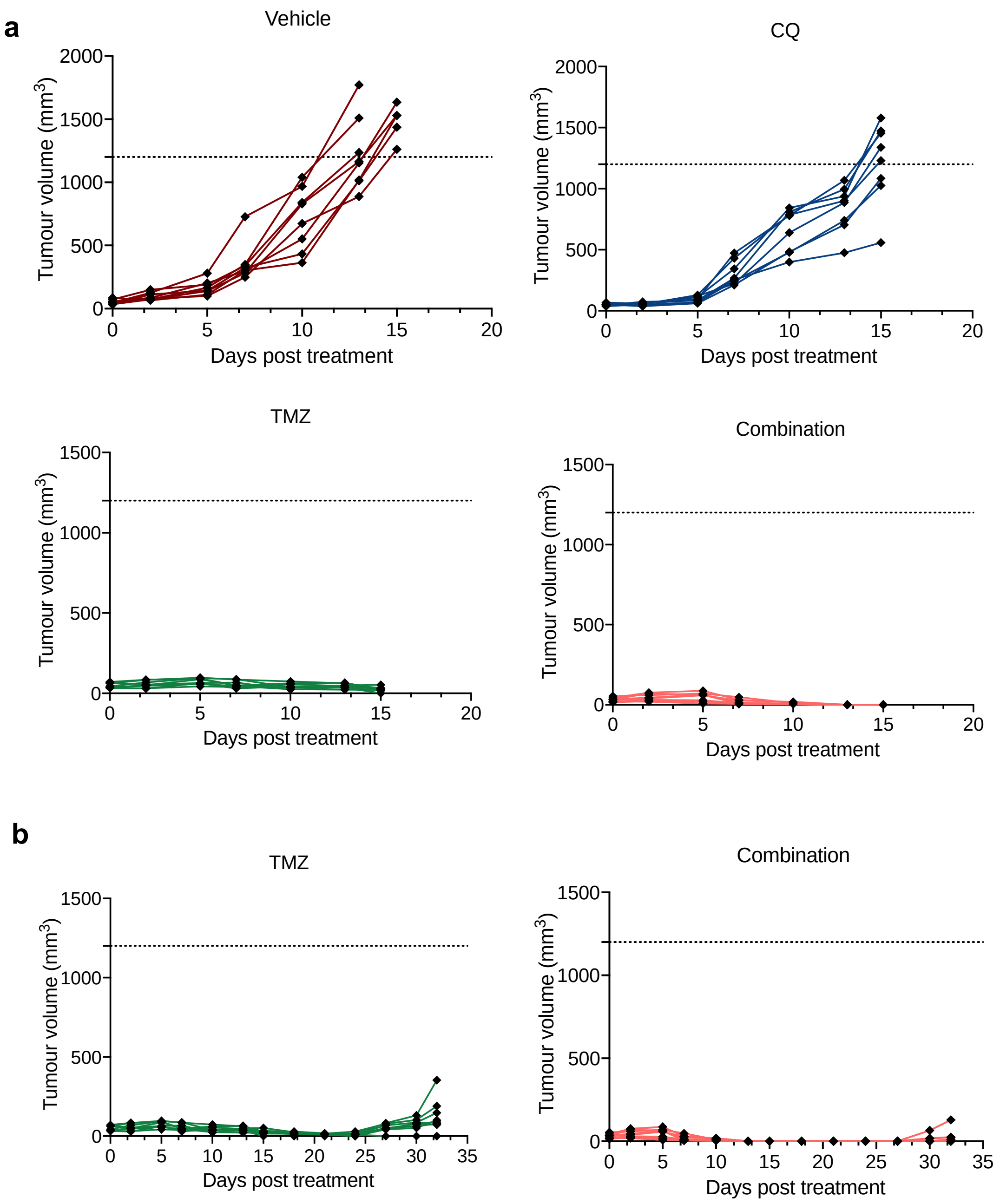


**a)** Individual tumor growth curve for each treatment group at day 15. Mice were culled if tumor volume reached 1200mm$^3$ (n=8). (**b)** Extended individual tumor growth for TMZ and combination group up to day 32 (n=8).

Supplementary Table S1: List of shRNA constructs

| Construct Name | Sequence |
|---|---|
| pSIH-H1-copGFP-shAIM2 #1 | CCAACTGGTCTAAGCAGCATT |
| pSIH-H1-copGFP-shAIM2 #2 | GCCACTAAGTCAAGCTGAAAT |
| pSIH-H1-copGFP-shAIM2 #3 | CTGGAGTTCATAGCACCATAA |
| pSIH-H1-copGFP-shGSDME #1 | GCATGATGAATGACCTGACTT |
| pSIH-H1-copGFP-shGSDME #2 | GCGGTCCTATTTGATGATGAA |
| pSIH-H1-copGFP-shGSDME #3 | GATGATGGAGTATCTGATCTT |
